# Characterization of a cytokinin-binding protein locus in *Mycobacterium tuberculosis*

**DOI:** 10.1101/2024.03.25.586654

**Authors:** Jin Hee Yoo, Cristina Santarossa, Audrey Thomas, Damian Ekiert, K. Heran Darwin

## Abstract

Cytokinins are adenine-based hormones that have been well-characterized in plants but are also made by bacteria, including the human-exclusive pathogen *Mycobacterium tuberculosis*. In *M. tuberculosis*, cytokinins activate transcription of an operon that affects the bacterial cell envelope. In plants, cytokinins are broken down by dedicated enzymes called cytokinin oxidases into adenine and various aldehydes. In proteasome degradation-deficient *M. tuberculosis*, the cytokinin-producing enzyme Log accumulates, resulting in the buildup of at least one cytokinin-associated aldehyde. We therefore hypothesized that *M. tuberculosis* encodes one or more cytokinin oxidases. Using a homology-based search for homologs of a plant cytokinin oxidase, we identified Rv3719 and a putative cytokinin-specific binding protein, Rv3718c. Deletion of the locus encoding these proteins did not have a measurable effect on *in vitro* growth. Nonetheless, Rv3718c bound a cytokinin with high specificity. Our data thus support a model whereby cytokinins play one or more roles in mycobacterial physiology.

**IMPORTANCE:** Numerous bacterial species encode cytokinin-producing enzymes, the functions of which are almost completely unknown. This work contributes new knowledge to the cytokinin field for bacteria, and also revealed further conservation of cytokinin-associated proteins between plants and prokaryotes.

## INTRODUCTION

*M. tuberculosis* is estimated to be present in almost one-third of the world’s population. Given that the existing vaccine is largely ineffective and that drug treatment is prolonged or otherwise ineffective in drug-resistant strains, there are ongoing efforts to identify new pathways to target for tuberculosis treatment. A major target of interest is the *M. tuberculosis* proteasome, which is essential to cause lethal infections in mice (1, 2). Proteasomes are highly regulated, multi-subunit, ATP-dependent proteases found in all domains of life, and it is known that *M. tuberculosis* proteasomes degrade numerous substrates with significant roles on bacterial physiology [reviewed in (3)]. Many proteins must first be post-translationally modified with the protein Pup that targets doomed proteins to an ATP-dependent hexameric ring called Mpa (mycobacterial proteasome ATPase; known as ARC in non-mycobacteria), which unfolds substrates to deliver them into a proteasome core protease. It was previously determined that the failed degradation of a single pupylated substrate, “lonely guy” (Log or Rv1205) results in bacteria that are highly sensitive to nitric oxide (NO) (4) and copper (Cu)(5).

Log was first characterized in plants and homologues are now well-recognized in numerous bacterial and fungal species. In both plants and *M. tuberculosis*, Log catalyzes the final biosynthetic step in cytokinin biosynthesis (4, 6). Cytokinins, which are adenine-based molecules with a modification at the nitrogen (*N*)^6^ position of the adenine rings, have been extensively studied in plants and are required for their normal growth and development (7, 8). In *M. tuberculosis,* cytokinins induce transcription of *loaA* (Rv0077c), resulting in a dramatic loss <u>o</u>f <u>a</u>cid-fast staining of bacteria (9). The mechanisms of how cytokinins induce *loaA* expression and how LoaA results in loss of acid-fast staining are unknown.

In plants, cytokinins are enzymatically degraded into adenine and aldehydes, presumably to abate signaling by these hormones as well as recycle their components; adenine is easily incorporated into numerous molecules while aldehydes likely react with other macromolecules or are otherwise detoxified. In an *mpa* mutant, the accumulation of Log results in high levels of several secreted cytokinins and at least one cytokinin-associated aldehyde called *para*-hydroxybenzaldehyde or *p*HBA, a breakdown product of *p*-topolin (10). Disruption of *log* is sufficient to reduce *p*HBA levels and fully restore NO and Cu resistance to an *mpa* mutant (4, 5). Importantly, *p*HBA is sufficient to sensitize wild-type (WT) *M. tuberculosis* to NO and Cu, suggesting this aldehyde directly disrupts resistance to these molecules (4, 5).

Enzymes involved in cytokinin breakdown in plants are called cytokinin oxidases or dehydrogenases (CKX), which remove chemical groups from the *N^6^* position of cytokinin adenine rings, liberating the associated aldehydes (10). We thus hypothesized that *M. tuberculosis* encodes a CKX needed to breakdown one or more cytokinins. We hypothesized disruption of a CKX-encoding gene in a proteasomal degradation-defective mutant would suppress the NO or Cu sensitive phenotypes of this strain, much like a *log* mutation (4, 5). In this work, we identified an open reading frame that encodes a protein with high similarity to a CKX from maize, but found that disrupting this gene did not suppress the NO or Cu sensitivity of an *mpa* mutant. However, we determined that Rv3718c, which is divergently expressed from Rv3719, encodes a cytokinin-binding protein similar to ones found in plants. Rv3718c bound to cytokinin with high specificity. Thus, our work has identified the first cytokinin-binding protein in bacteria.

## RESULTS AND DISCUSSION

### *M. tuberculosis* encodes putative cytokinin-associated proteins

We tested the hypothesis that *M. tuberculosis* has a CKX protein by performing a BLASTP search (11) using *Zea mays* (maize, corn) cytokinin dehydrogenase 1 precursor (accession number BAA74451) against the *M. tuberculosis* proteome. The protein with the highest identity (82/212, 39%) was Rv3719 (**Fig. 1A**). Despite numerous efforts including different epitope tags and bacterial expression systems, we could not produce recombinant Rv3719. Using ProtParam (12), we found the instability index for Rv3719 is 48.04 (unstable), which supports our inability to produce this protein.

**Fig. 1.**
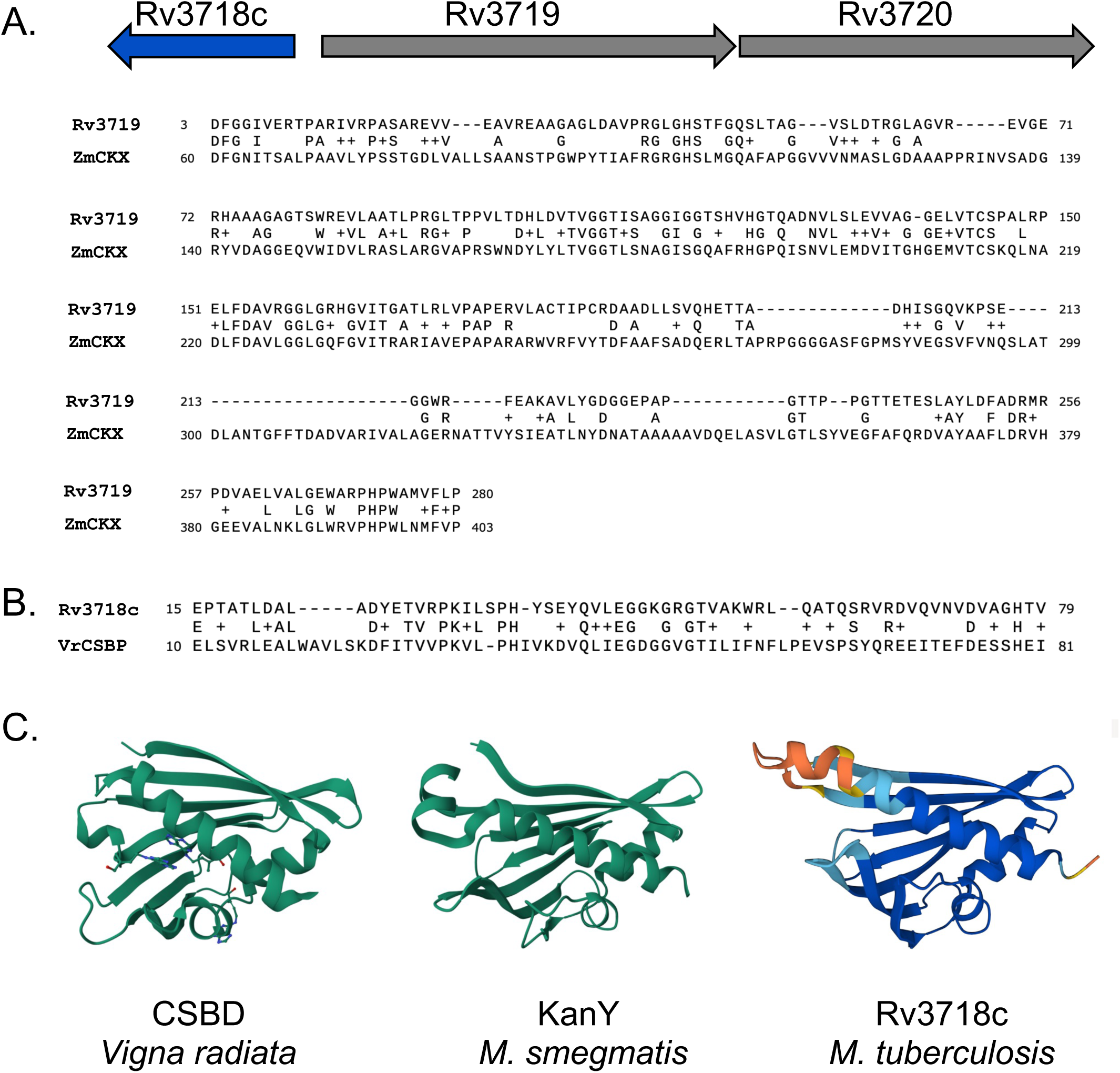
Putative cytokinin-binding proteins in *M. tuberculosis*. **(A)** Alignment of *Zea mays* CKX with *M. tuberculosis* H37Rv Rv3719. **(B)** Alignment of *Vigna radiata* (Vr) CSBP with Rv3718c. **(C)** Comparison of VrCBSP (PDB: 2FLH) with *M. smegmatis* KanY (PDB: 5WOX) and the AlphaFold model of Rv3718c. Molecules in VrCSBP are the cytokinin zeatin.

Downstream of Rv3719 is Rv3720, which is predicted to encode a fatty acid synthase. Given that some cytokinins have prenyl groups, it is possible Rv3720 binds to one or more of these hydrophobic molecules for their metabolism. Divergently expressed from Rv3719 is Rv3718c, which has also been called *kanY* in *M. smegmatis* (GenBank: STZ35122.1) (**Fig. 1B**). Interestingly, all three genes are conserved in *M. leprae*, which has a highly decayed genome (13). This observation suggests it plays a core function in mycobacterial physiology. Comparison of either the nuclear magnetic resonance (NMR) structure of MSMEG_6282 or an AlphaFold model of Rv3718c (14) to the cytokinin-specific binding protein (CSBP) in *Vigna radiata* (mung bean) revealed strong structural similarities among the proteins (**Fig. 1C**).

### Deletion of the Rv3718c-Rv3720 locus did not alleviate the NO sensitivity of a proteasomal degradation-defective strain

We hypothesized that the disruption of a putative CKX would reduce the production of aldehydes from cytokinin breakdown in a proteasomal degradation mutant. For simplicity, we simultaneously deleted and disrupted all three genes from the *M. tuberculosis* chromosome with the expectation that their complete removal would increase our chances of having an observable phenotype. We deleted these genes from both the WT and *mpa* strains of *M. tuberculosis* and confirmed the loss of Rv3718c by immunoblotting (**Fig. 2A**). Deletion of this locus had no measurable impact on growth under routine culture conditions (**Fig. 2B**). We next tested whether or not deletion of these genes restored NO resistance to a proteasomal degradation-defective strain. Unlike disruption of *log* (4), deletion of this locus did not suppress the NO-sensitive phenotype of the *mpa* mutant (**Fig. 2C**).

**Fig. 2.**
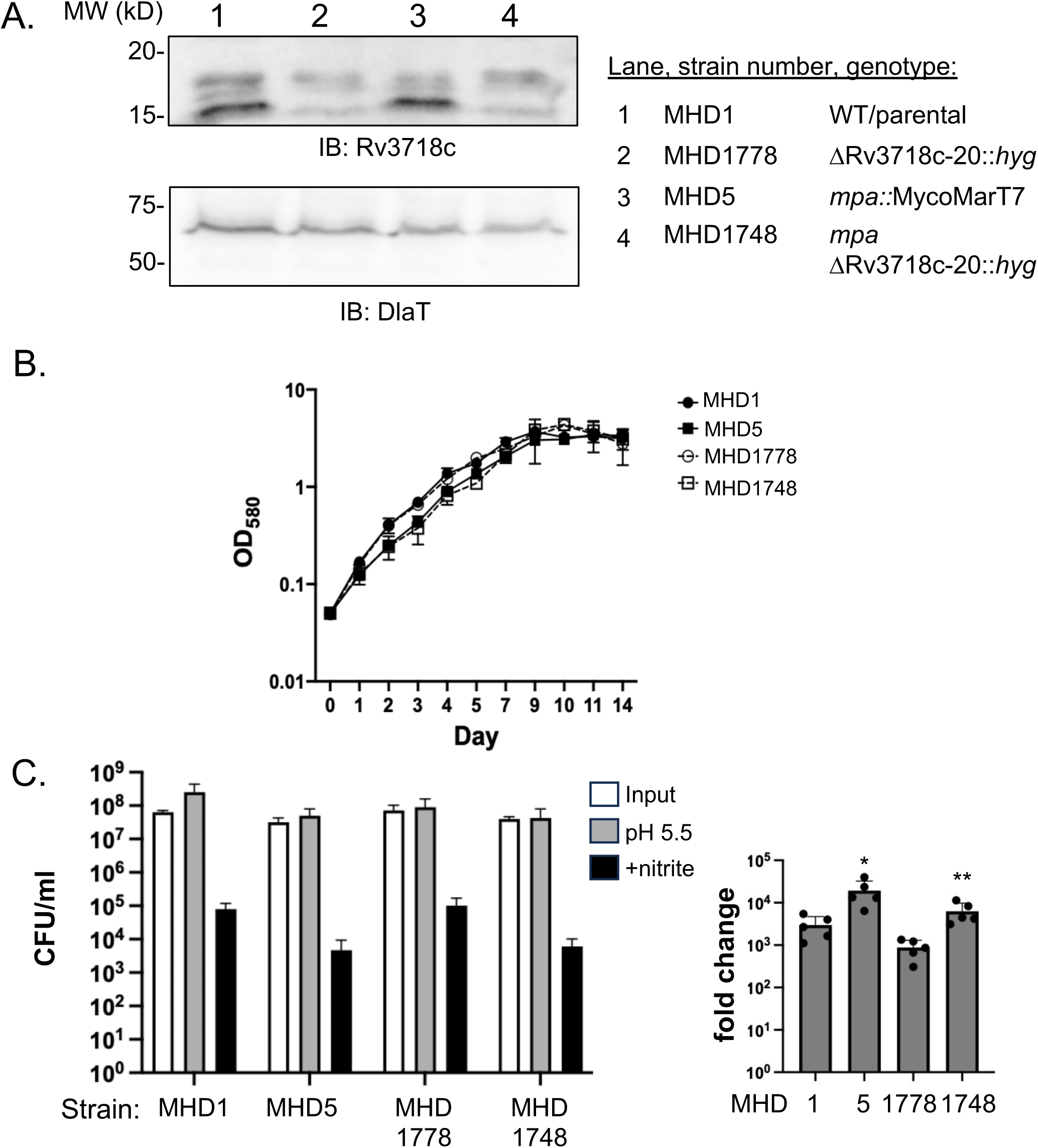
Deletion of Rv3718c-Rv3720 locus did not affect growth under routine culture conditions or resistance to nitric oxide. **(A)** Rv3718c was undetectable in strains lacking the Rv3718c-Rv3720 locus. IB = immunoblot. DlaT (dihydrolipoamideacyltransferase) served as a loading control. **(B)** Deletion of Rv3718c-Rv3720 did not affect growth in Middlebrook 7H9c broth. Growth curves were performed in duplicate and are representative of two independent experiments. **(C)** Deletion of Rv3718c-Rv3720 did not affect sensitivity to nitric oxide (NO). NO assays were performed in technical triplicate and are representative of two independent experiments with an outlier technical replicate omitted. Given the significant difference in growth between the parental and mpa strains without NO, we calculated the fold change in survival in NO compared to acidic media alone, which more accurately represents the susceptibility of the strains to NO. * indicates *P* = 0.02, ** indicates *P* = 0.008.

### Rv3718c is a cytokinin-specific binding protein

To test if Rv3718c could bind a cytokinin or related molecules, we used microscale thermophoresis (MST). We purified C-terminally, hexahistidine-tagged Rv3718c from *E. coli* (**Fig. 3A**) see Materials and Methods and Table 1) and found Rv3718c bound the cytokinin N^6^-(2-isopentenyl)adenine (2iP) with an estimated K_d_ of ∼125-250 µM (**Fig. 3B**). At the concentrations tested, Rv3718c did not bind *N*^6^-(Δ^2^-Isopentenyl)adenosine (iPR), the ribose precursor of 2iP, or adenine, a precursor and breakdown product of 2iP (**Fig. 3, far right**). These data suggest Rv3718c is a cytokinin-specific binding protein. Given the structural and functional similarities with plant CSBP, we suggest naming the mycobacterial protein Cbp for “<u>c</u>ytokinin-specific <u>b</u>inding <u>p</u>rotein”.

**Fig. 3.**
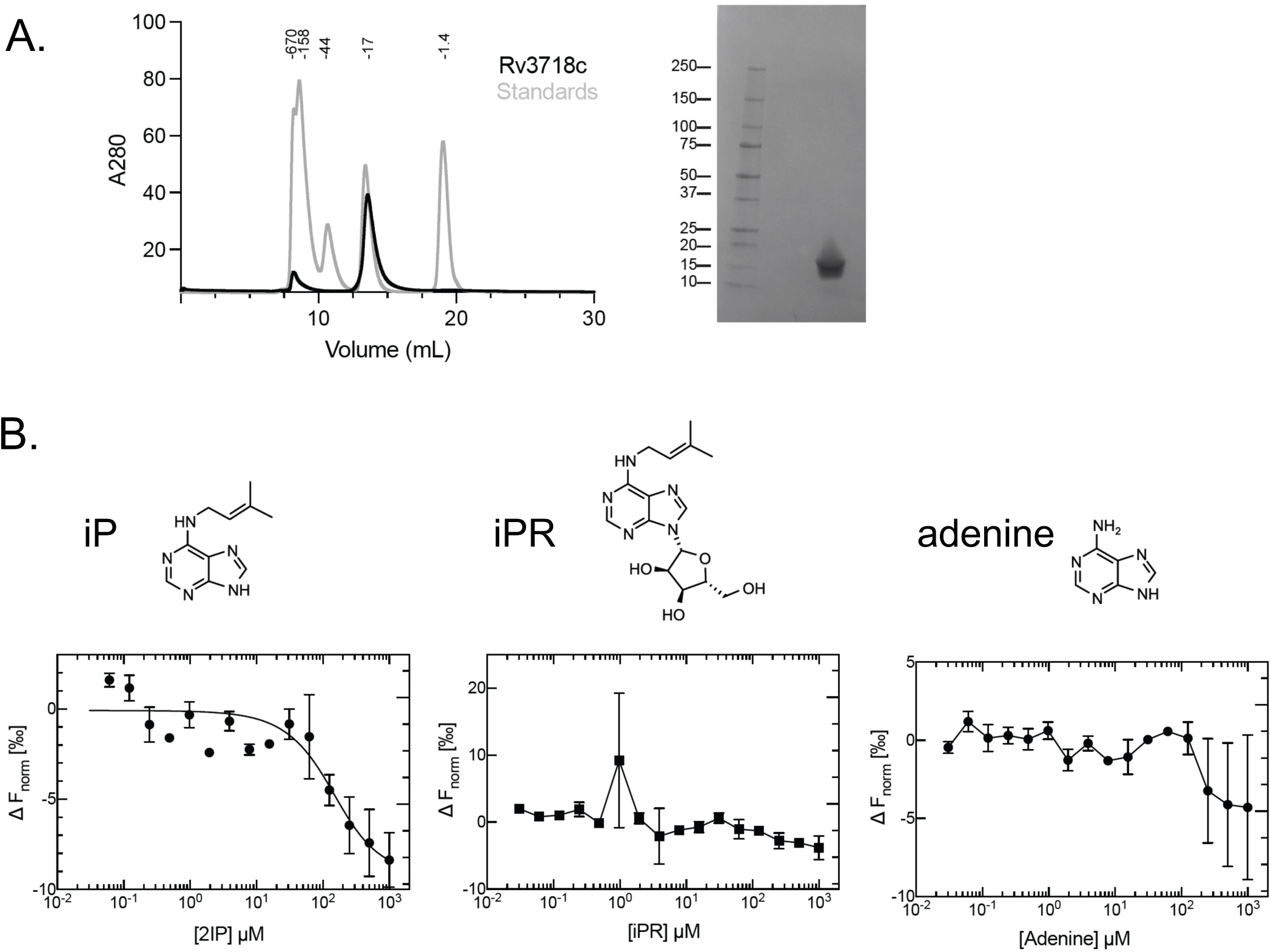
Rv3718c is a cytokinin-specific binding protein. Purification of Rv3718 and characterization of cytokinin binding. **(A)** Right: Size exclusion chromatography of Rv3718c-His6 (black) overlaid with molecular weight standards (gray). Left: SDS-PAGE gel of purified Rv3718c-His6. **(B)** Microscale thermophoresis (MST) analysis to assess binding of the indicated compounds (above) to Rv3718c-His_6_. Data are shown as mean ± range and were fit using Equation 1. The individual data points are shown from two independent experiments.

**Table 1.**
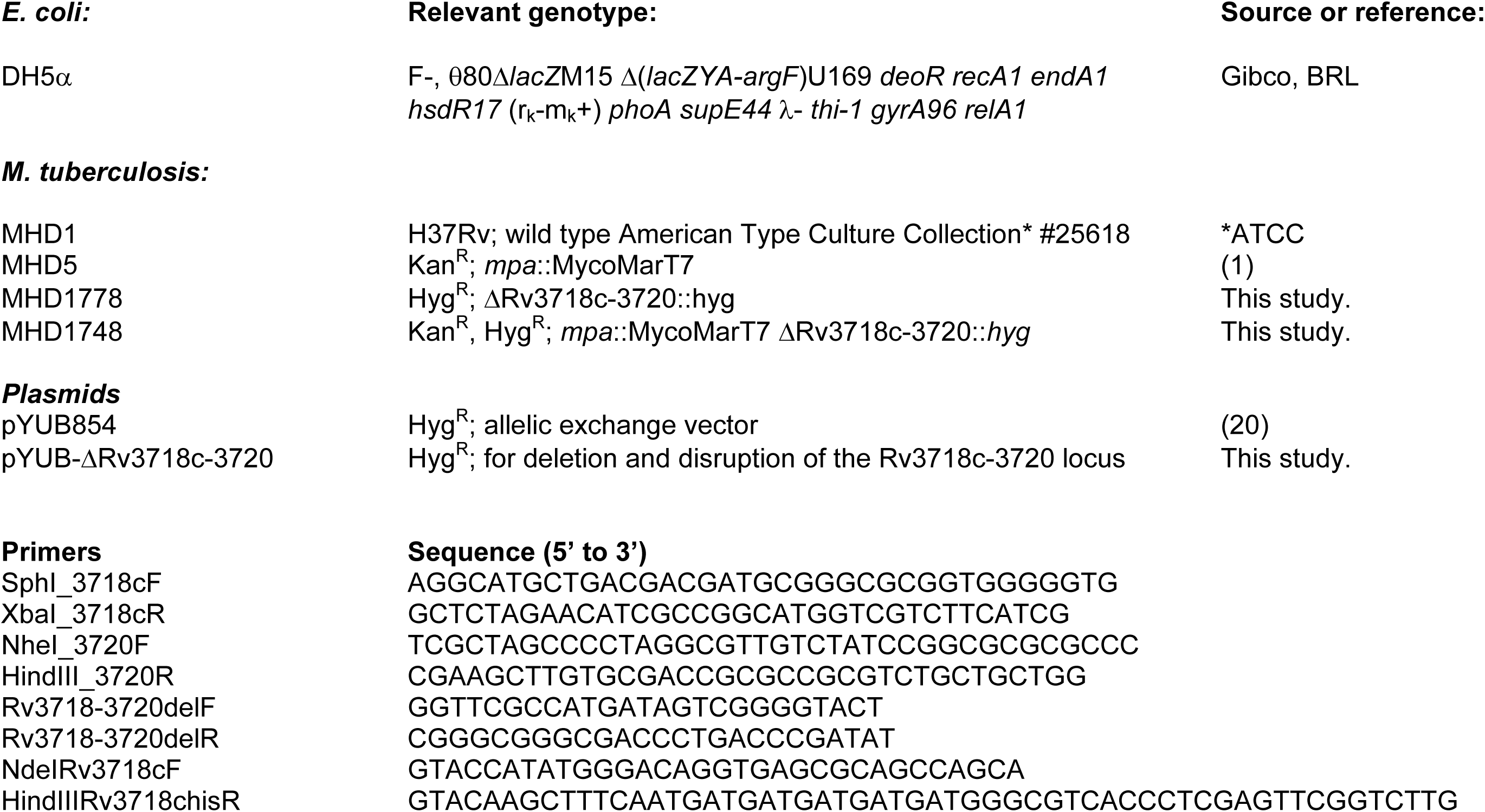
Bacterial strains, plasmids and primers used in this work.

While the failed degradation of the cytokinin activating enzyme Log weakens *M. tuberculosis* during infections (4), the genes in the *cbp*-Rv3719-20 locus are not essential for *M. tuberculosis* pathogenesis in at least one infection model (15). Nonetheless, the conservation of cytokinin signaling in mycobacteria, including *M. leprae*, strongly suggests it plays an important function in physiology at some specific point of a *Mycobacterium* life cycle, for example, specific to human or other infections. In addition to understanding the function of cytokinins in bacterial physiology, it remains to be determined how cytokinins transduce signals to affect gene expression. Plants use two-component regulatory systems (TCSs) closely related to bacterial TCSs. Although *M. tuberculosis* encodes at least 16 TCSs, the only transcription factor that is linked to cytokinin-inducible gene expression is a TetR family repressor called LoaR (Rv0078c). LoaR does not bind cytokinin and its native ligand has not been identified (9). Finally, because plant CBP activity has not been determined, it is possible the characterization of bacterial Cbps might help one day elucidate the role of these proteins in plant biology.

## MATERIALS AND METHODS

### Strains, plasmids, primers, and culture conditions

See Table 1 for strains, plasmids, and primers used in this work. Reagents used for making all buffers and bacterial media were purchased from ThermoFisher Scientific, unless otherwise indicated. *M. tuberculosis* was grown in "7H9c" [BD Difco Middlebrook 7H9 broth with 0.2% glycerol and supplemented with 0.5% bovine serum albumin (BSA), 0.2% dextrose, 0.085% sodium chloride, and 0.05% Tween-80]. For solid media, *M. tuberculosis* was grown on Middlebrook 7H11 agar (“7H11”, BD Difco) containing 0.5% glycerol and supplemented with 10% final volume of BBL Middlebrook OADC Enrichment. For selection of *M. tuberculosis*, the following antibiotics were used as needed: kanamycin 50 µg/ml, hygromycin 50 µg/ml. *E. coli* was cultured in Luria-Bertani (LB) broth or on LB agar (both BD Difco). Media were supplemented with the following antibiotics as needed: kanamycin 100 µg/ml and hygromycin 150 µg/ml. Cytokinin (N^6^-(2-isopentenyl)adenine was purchased from Sigma Aldrich (Catalog #D7674), adenine was purchased from Sigma (Catalog #A8626), and N^6^-(Δ^2^-isopentenyl)adenosine was purchased from Cayman Chemical Company (Catalog #20522).

To make the pYUBΔRv3718c-Rv3720 allelic exchange plasmid, 700bp upstream of Rv3718c and downstream of Rv3720 were amplified by PCR and cloned into pYUB854 flanking a hygromycin resistance cassette (see Table 1 for primers). For pET24b(+)-Rv3718c, primers NdeIRv3718cF and HindIIIRv3718chisR were used to PCR amplify Rv3718c from genomic DNA. *E.coli* DH5α was used for transformations. Plasmids were purified from *E. coli* using the QIAprep Spin Miniprep Kit (Qiagen). All plasmids made by PCR cloning were sequenced by Azenta, Inc. to ensure the veracity of the cloned sequence. Primers used for PCR amplification or sequencing were purchased from Life Technologies or IDT and are listed in Table 1. DNA was PCR-amplified using Phusion polymerase (New England Biolabs; NEB) according to the manufacturer’s instructions. PCR products were purified using the QIAquick Gel Extraction Kit (Qiagen). Restriction enzymes and T4 DNA ligase were purchased from NEB. *M. tuberculosis* was transformed by electroporation as previously described (16).

For allelic exchange we used a previously described method (17). Chromosomal DNA was purified from transformants and screened by PCR using primers Rv3718-3720delF and Rv3718-3720delR to determine if the locus was lost from transformants.

All *M. tuberculosis* work was performed in the ABSL3 facility of NYU Grossman School of Medicine, in accordance with its Biosafety Manual and Standard Operating Procedures.

### Purification and labeling of Rv3718c

Hexahistidine-tagged Rv3718c was produced in *Escherichia coli* BL21(DE3) cells and grown in LB at 37°C. Expression was induced at OD_600_ 0.6-0.7 with 0.1 mM IPTG and grown for four hours at 37°C. The cultures were pelleted and resuspended in buffer containing 50 mM NaH_2_PO_4_ pH 8.0, 100 mM NaCl, 10 mM imidazole. All subsequent purification steps were performed at 4°C. Cells were lysed using the Emulsiflex-C5 homogenizer (Avestin, 2 cycles at 15,000 psi) and the homogenized lysate was centrifuged at 30,000 *g* for 1 hour. The clarified lysate was applied onto a column containing Ni Sepharose 6 resin (Cytiva Life Sciences) and washed with 20 column volumes of buffer containing 50 mM NaH_2_PO_4_ pH 8.0, 100 mM NaCl, 20 mM imidazole. The protein was eluted with buffer containing 50 mM NaH_2_PO_4_ pH 8.0, 100 mM NaCl, 250 mM imidazole. The eluent was concentrated using an Amicon Ultra 3 kDa MWCO centrifugal filter and then exchanged into size-exclusion buffer (20 mM HEPES pH 8.0, 100 mM NaCl, 5% glycerol) using Zeba™ Spin Desalting Columns (MWCO 7 kDa, 0.5 mL). The protein was labeled with Rv3718c was labeled with Alexa Fluor 647 NHS Ester (Invitrogen™ Catalog# A20106) for 30 minutes at room temperature in the dark. The sample was loaded onto a Superdex™ 75 Increase 10/300 GL column (Cytiva Life Sciences) in size-exclusion buffer. Fractions were pooled, yielding protein concentrations of 8-14 µM, flash frozen in liquid nitrogen and stored at - 80°C.

### Microscale thermophoresis assay (MST)

The binding affinities of 2iP, iP and iPR to Rv3718c were measured using Monolith NT.115 (NanoTemper Technologies, Munich, Germany). Frozen protein aliquots were thawed and diluted in MST buffer (20 mM HEPES pH 8.0, 100 mM NaCl, and 5% glycerol, 0.005% Tween 20). To obtain a fluorescence intensity between 200 and 1100 counts at 20% LED power, a final concentration of 100 nM protein was used. A 25X compound stock was prepared by a 1:1 serial dilution of the compound in DMSO starting from 25 mM and then diluted in MST buffer to obtain a 2X compound stock. The 2X protein stock (200 nM Rv3718c) was mixed well with 2X compound and incubated for 5 min at RT in the dark. The final DMSO concentration is 4%. Each sample was transferred to a Monolith premium capillary tube, and measurements were performed at 20, 40 and 60% MST power at 25°C. At 60% MST power, a binding curve for 2iP was observed at 10s after the start of thermophoresis. The ⊗F_norm_ values were fitted to Equation (1) using Prism v. 10.1.1 (GraphPad Software Inc) at different concentrations of 2iP.

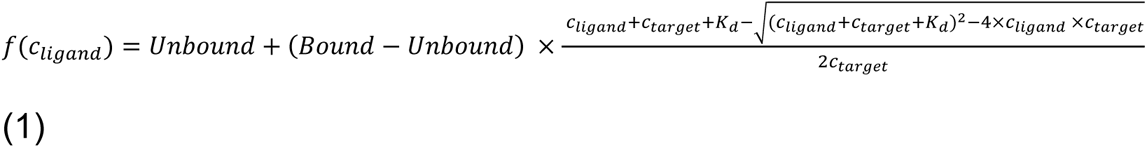

In Equation (1), *f(c_ligand_)* is the F_norm_ value at a given ligand concentration *c_ligand_*, *Unbound* is the F_norm_ signal of the target alone, *Bound* is the F_norm_ signal of the complex, *K_d_* is the dissociation constant or binding affinity, and *c_target_* is the final concentration of the target.

### Preparation of *M. tuberculosis* extracts for immunoblotting

*M. tuberculosis* cultures were grown to an optical density at 580 nm (OD_580_) of ∼1. Equivalent cell numbers were collected based on the OD_580_ of the cultures. For example, an “OD_580_ equivalent of 1” indicates the OD_580_ of a 1 ml culture is 1.0. Five OD_580_ equivalents of bacteria were harvested by centrifugation at 3,000 *g,* washed in PBST (PBS, 0.05% Tween 80), resuspended in lysis buffer (100mM Tris-Cl pH8, 1mM EDTA pH8), and transferred to a tube containing 250 µl of 0.1 mm zirconia beads (BioSpec Products). Bacteria were lysed using a mechanical bead-beater (BioSpec Products). Whole lysates were mixed with 4· reducing SDS sample buffer (250 mM Tris pH 6.8, 2% SDS, 20% 2-mercaptoethanol, 40% glycerol, 1% bromophenol blue) to a 1· final concentration, and samples were boiled for 10 minutes at 100°C.

For immunoblotting, whole cell lysates were separated by 10% sodium dodecyl sulfate-polyacrylamide gel electrophoresis (SDS-PAGE). Proteins were transferred to nitrocellulose membranes (GE Amersham) by semi-dry transfer for 10 min at 15V (Bio-Rad), air dried, then blocked with 3% BSA in TBST. Horse-radish peroxidase conjugated secondary antibodies to rabbit IgG were purchased from Pierce. Immunoblots were developed using SuperSignal West Pico PLUS chemiluminescent substrate (ThermoFisher Scientific) and imaged using Bio-Rad ChemiDoc system and quantified using Fiji (18). Blots were stripped as previously described (19). For loading control, the membrane was reblocked with 2% milk in 1· TBST, incubated with polyclonal antibodies to DlaT (kind gift from R. Bryk and C. Nathan), and imaged as above.

### Nitric oxide sensitivity assay

Assays were performed as described previously (1). Briefly, bacteria were grown to an OD_580_ ∼0.8-1 and resuspended in acidified 7H9c (pH 5.5) and diluted to an OD_580_ of 0.08. Bacteria were then aliquoted in triplicate to flat-bottom 96-well plates and a fresh sodium nitrite solution was added to each well at a final concentration of 3 mM. Bacteria were incubated for six days at 37°C before plating onto 7H11 OADC Y-plates and incubated at 37°C for enumeration 2-3 weeks later. Data were graphed and analyzed in Prism.

## Acknowledgments

We thank P. Tran and G. Bhabha for reading a draft version of this manuscript. We thank A. Jordan for plasmid construction (supported in part by a Public Health Service Institutional Research Training Award T32 AI007180). This work was supported by NIH grants AI088075 and AI153197 awarded to K.H.D. D.E. was supported by NIH grant AI174646. We thank the Office of Science & Research High-Containment Laboratories at NYU Grossman School of Medicine for their support in the completion of this research.

## Author contributions

C.S., J.H.Y., and K.H.D. designed the research; C.S., J.H.Y., and A.T. performed research; C.S., J.H.Y., D.E., and K.H.D wrote the manuscript. The authors declare that they have no conflicts of interest with the contents of this article.

## DATA AVAILABILITY

All data supporting these findings are available within the article and/or its supplementary materials.

